# Integrative single-cell and spatial mapping of oxidative stress response uncovers GCLC+ mesenchymal tumor cell state linked with favorable outcomes in triple negative breast cancer

**DOI:** 10.1101/2025.11.06.686771

**Authors:** Nomeda Girnius, Tuulia Vallius, Wenqing Chen, Inga-Maria Launonen, Sara Palomino-Echeverria, Jia-Ren Lin, Caitlin E. Mills, Silja Kauppila, Pauliina Kronqvist, Antti Ellonen, Merja Perala, Eloise Withnell, Yu-An Chen, Maria Secrier, Sandro Santagata, Peter K. Sorger, Anniina Farkkila

## Abstract

Inhibiting oxidative stress response (OSR) proteins has been suggested as a therapeutic strategy in triple negative breast cancer (TNBC). However, the cell type specificity and spatial distribution of OSR genes and proteins, such as GCLC and NQO1, is unknown. Using single cell and spatial transcriptomics datasets we found that OSR genes were highly expressed in TNBC tumor cells, which localized in spatial clusters. Multiplex immunofluorescence imaging of 345 TNBC samples from 186 patients demonstrated that OSR proteins GCLC and NQO1 exhibit distinct expression profiles across tumor, immune, and stromal cell populations and are elevated in inflamed histological regions. Tumor cell OSR protein expression was associated with the composition of the adjacent cellular neighborhood. Furthermore, we identified GCLC and vimentin positive (GCLC+VIM+) mesenchymal-like tumor cells, residing near immune cells and exhibiting increased proliferation and decreased anastasis signatures, suggesting sensitivity to chemotherapy. Across a panel of thirteen TNBC cell lines, GCLC expression was positively correlated with sensitivity to cisplatin. Cox regression analysis revealed that patients with higher proportions of GCLC+VIM+ tumor cells had a longer overall survival. Collectively, our results demonstrate that individual OSR proteins are expressed in distinct microenvironments and tumor cell states, potentially contributing to patient outcomes.

## Introduction

Triple negative breast cancer (TNBC) is a highly invasive disease that accounts for up to 15% of breast cancers, has a poor patient prognosis, and limited treatment options^1–4^, involving chemotherapy, surgery and radiation, with the recent addition of immune checkpoint inhibitors^5^. Recent studies in TNBC have demonstrated a relationship between the spatial tumor microenvironment (TME) and disease progression and patient outcomes^6–9^, highlighting the importance of understanding cellular organization within the tumor. New therapies based on the biology of TNBC are sorely needed.

A potential target in TNBC could be oxidative stress response (OSR) proteins, as they can be transcriptionally upregulated in this disease, suggesting elevated oxidative stress^10^ might be a viable therapeutic opportunity^11–14^. In the presence of elevated oxidative stress, the master regulator of the antioxidative response NRF2 (encoded by *NFE2L2*) is stabilized leading to expression of a host of genes from different pathways representing multiple mechanisms of defense against oxidative damage^15^. Elevated expression of certain NRF2 targets, for example NAD(P)H quinone dehydrogenase 1 (NQO1)^16^, thioredoxin reductase 1 (TXNRD1)^17^, and glutamate-cysteine ligase catalytic subunit (GCLC)^18^ has been reported in breast cancer and associated with worse outcomes. A detailed characterization of OSR gene expression in the context of the diverse TNBC tumor cell phenotypes has not been performed, and the expression of these proteins in the immune and stromal compartments has not been described.

Using single-cell RNA sequencing (scRNAseq) and spatial transcriptomics data together with cyclic immunofluorescence data from over 180 patients, we explored the phenotypes and spatial distribution of TNBC tumor cells expressing OSR genes and proteins. We found that diverse cell types engage different antioxidative responses, with NQO1 being expressed in tumor cells, and GCLC mainly in stromal and immune populations. Through single-cell profiling, we identified a distinct tumor cell population characterized by co-expression of GCLC and the mesenchymal marker, vimentin (VIM). These cells were found to spatially colocalize with multiple immune cell types and were associated with better patient outcomes. In addition, cell line data suggest that GCLC+ VIM+ tumor cells may also be more susceptible to first line chemotherapies used in TNBC. This study provides new insights into OSR protein expression in TBNC and provides a framework for studying OSR genes and proteins, and understanding the consequences of targeting them therapeutically.

## Results

### Single-cell and spatial transcriptomics reveal high oxidative stress response (OSR) gene expression in TNBC

We studied OSR in TNBC using a multimodal approach (Fig. 1a), first focusing on publicly available transcriptomics data. To determine the cell type specificity and patterns of OSR gene expression in TNBC, we developed a custom OSR signature consisting of genes encoding proteins involved in the antioxidative response (hereafter referred to as AntiOx signature; Supplementary Table 1). In publicly available breast cancer scRNASeq data^19^ (Fig. 1b), TNBC tumor cells had a significantly higher expression of the AntiOx signature relative to ER+ tumor cells (Fig. 1c), with highest expression in the tumor cells followed by myeloid, perivascular-like cells (PVL), and cancer-associated fibroblasts (CAFs) (Supplementary Fig. 1a). We observed a high degree of inter- and intratumoral heterogeneity in the enrichment of the AntiOx signature in tumor cells across patients (Fig. 1d, Supplementary Fig. 1b). While a block of genes were highly correlated, many other genes showed weaker correlations suggesting a less coordinated gene expression program (Supplementary Fig. 1c).

**Figure 1.**
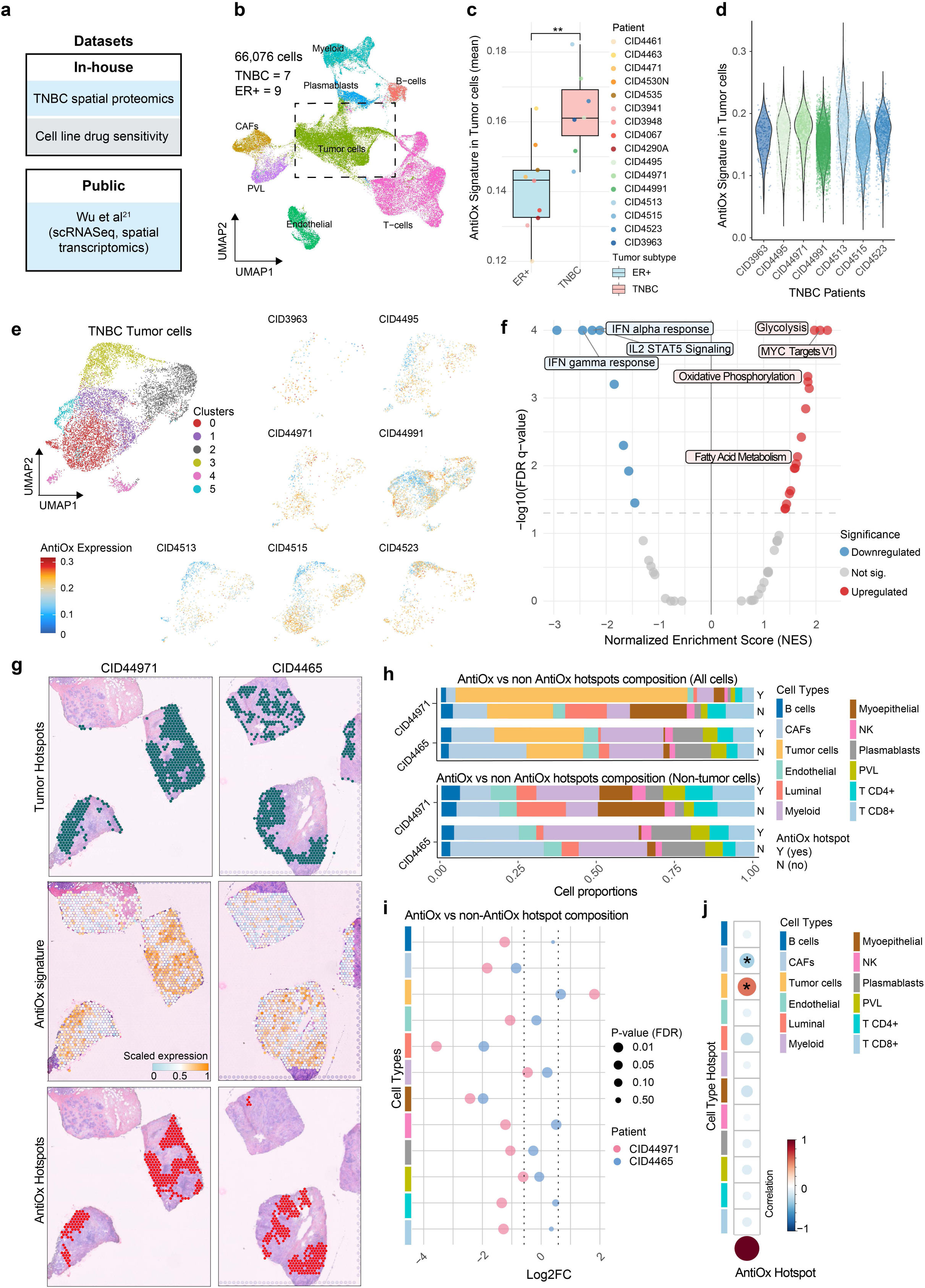
Distinct single-cell and spatial OSR gene expression in TNBC. **a** Summary of datasets used in manuscript. **b** Uniform manifold approximation and projection (UMAP) of 66,076 single cells annotated for cell type, reanalyzed from Swarbrick et al^19^. **c** Boxplot of the AntiOx signature expression in ER+ breast cancer and TNBC. The box represents the interquartile range (IQR) from the first quartile (Q1) to the third quartile (Q3), with the horizontal line indicating the median. Whiskers extend to 1.5 × IQR from the quartiles. Asterisks indicate *p < 0.01* based on the unpaired Wilcoxon rank-sum test. **d** Violin plots showing the AntiOx signature scores across TNBC samples. **e** UMAPs showing AntiOx signature expression across tumor cell subclusters of TNBC. **f** Volcano plot showing Hallmark pathways that are significantly upregulated (right side) or significantly downregulated (left side) in AntiOx^High^ tumor cells compared to AntiOx^Low^ tumor cells. **g** An H&E of Visium data is presented together with coloring to show tumor cell-rich spots (top), AntiOx signature expression in the spots (middle), and the identified AntiOx hot spots (bottom). **h** Stacked bar plot summarizing cell type composition in AntiOx hot spots and non-hotspots in all cells (upper panel) and rescaled without tumor cells (lower panel). **i** Cleveland plot showing the log2 fold-change in hotspot composition between AntiOx and non-AntiOx conditions for each cell type. Hotspots were considered differentially expressed if the log2 fold-change was greater than 0.58 (indicated by dashed vertical lines) and the adjusted p-value was less than 0.05 (reflected by dot size). **j** Correlation of cell type hotspots with AntiOx hotspots. Higher color intensity and larger dot size indicate stronger correlations. Positive and negative correlations are shown in red and blue, respectively. Asterisks indicate statistically significant correlations.

To assess whether high OSR gene expression is associated with a specific tumor cell subpopulation, we performed unsupervised clustering of TNBC tumor cells which resulted in six distinct tumor cell clusters (Fig. 1e). The AntiOx signature was expressed similarly across clusters suggesting that different cell states may be associated with the antioxidative response. The clusters with the highest AntiOx signature were 0, 2, and 4. Both clusters 0 and 4 were characterized by immune-related pathways, for example those involved in antigen presentation and immune response. Cluster 2 highly expressed pathways associated with cell division (Supplementary Fig. 1d-f, Supplementary Table 2).

Elevated oxidative stress and increased OSR gene expression have been linked to sustained proliferation and increased metabolic activity^20^, inflammation in the microenvironment^21^, and growth outside of a normal cellular niche^22^. To explore the association between metabolism, proliferation, and inflammation and elevated OSR gene expression in human tissues, we compared the AntiOx signature-high (top 25%, AntiOx^High^) and AntiOx signature-low (bottom 25%, AntiOx^Low^) tumor cells by performing differential gene expression analysis followed by gene set enrichment analysis (Fig. 1f; Supplementary Table 3). The AntiOx^Low^ cells were enriched for pathways associated with inflammation (Allograft Rejection, IFNa and IFNy, TNFa, IL6-JAK-STAT3, Inflammatory Response, IL2-STAT5, and Complement). In contrast, AntiOx^High^ tumor cells were enriched for cell cycle (E2F Targets, MYC Targets v1 and v2, G2M Checkpoint) and metabolic (Glycolysis, Oxidative Phosphorylation, Fatty Acid Metabolism, Adipogenesis, MTORC1 signaling) pathways, indicative of cell proliferation-related biological processes. This suggests that tumor cell-intrinsic inflammatory signaling is not necessarily needed for the induction of the OSR gene expression in tumor cells.

To determine the contribution of the TME and cellular niches to OSR gene expression, we used a Visium spatial transcriptomics dataset^19^ that shared samples with the scRNASeq data set (slide 1: CID44971; slide 2: CID4465) and probed the spatial distribution of the AntiOx signature. Using a spatial statistics tool SpottedPy^23^, we identified “hotspots,” spatial clusters with significantly enriched gene expression scores (e.g., AntiOx or tumor cell signatures). This approach provides a systematic way to delineate statistically significant hotspots in spatial transcriptomics data, allowing for quantification of their extent and significance. We identified significant AntiOx hotspots which visually colocalized with tumor spots (Fig. 1g). Visium data contains on average 15–20 cells per spot, which can be deconvolved using the Cell2location method^24^.This method also showed that most of the AntiOx hotspots were predominantly composed of tumor cells (Fig. 1h, top), followed by cancer-associated fibroblasts (CAFs) and myeloid cells (Fig. 1g, bottom). Luminal cells, a subset of immune cells (CD8+ T cells, CD4+ T cells, B cell, NK cells) and other stromal cells (PVL, endothelial cells) were also detected but accounted for a minority of the cells in the AntiOx hotspots. A comparison between AntiOx hotspots and the remaining spots revealed consistently increased tumor cells and decreased cancer-associated fibroblasts (CAFs), myoepithelial and luminal cells in the hotspots (Fig. 1i; see also Supplementary Fig. 1g). To further investigate the association of the AntiOx signature with TME cell types, we identified hotspots for the twelve major deconvoluted cell types (see Methods for details). The correlation of the different cell type hotspots with the AntiOx hotspots confirmed a significant positive correlation between tumor cells and the AntiOx signature (Fig. 1j).

Collectively, our data show that OSR genes are expressed in tumor cells as well as myeloid cells and fibroblasts, highlighting that additional cell types express OSR genes in tumors. Furthermore, the AntiOx signature shows a non-random distribution, most often co-localizing with the tumor-rich areas.

### Multiplex imaging identifies a unique tumor cell population characterized by GCLC and Vimentin expression

To further assess OSR protein spatial distribution in patient samples, and to investigate the association of the microenvironment with OSR protein expression, we performed cyclic immunofluorescence (CyCIF) imaging of a large cohort of primary TNBC tumors (345 cores from 186 patients) using a tissue microarray (TMA) (Fig. 2a). We collected detailed clinical information for each patient including tumor stage, patient age, and overall survival (Fig. 2b; Supplementary Table 4). Importantly, the TMA was constructed using larger than usual 1.5 mm cores (compared to 1 mm) containing a mean of 10671.8 cells per core and multiple histological regions within a single tumor were sampled thus enabling intra-tumor comparisons of tumor center (n=192), invasive border (n=66), inflamed regions (n=61), and lymph node metastases (n=26) (Supplementary Fig. 2a-b). We used a 34-marker antibody panel focused on TME cell type identification, assessment of signaling pathway activity, and OSR protein expression (Supplementary Table 5). Because the OSR has distinct mechanisms for detoxification, we selected markers that would represent three of these mechanisms to broadly capture the OSR program.We included validated antibodies against NQO1 for quinone detoxification, GCLC for the glutathione pathway, and TXNRD1 for the thioredoxin pathway. All three were also highly expressed in AntiOx^High^ cells (Supplementary Fig. 2c) Over 3.5 million high-quality single cells passed quality control (see Methods) for downstream spatial analyses. We identified tumor (panCK+), immune (CD8+ T cells, CD4+ T cells, regulatory T cells (Foxp3+ CD4+), CD68+ macrophages, CD206+ macrophages, and other CD45+ immune cells) and stromal (SMA+, VIM+, and other panCK-CD45-cells) cell populations, which occurred at different proportions across the patients (Supplementary Fig. 2d).

**Figure 2.**
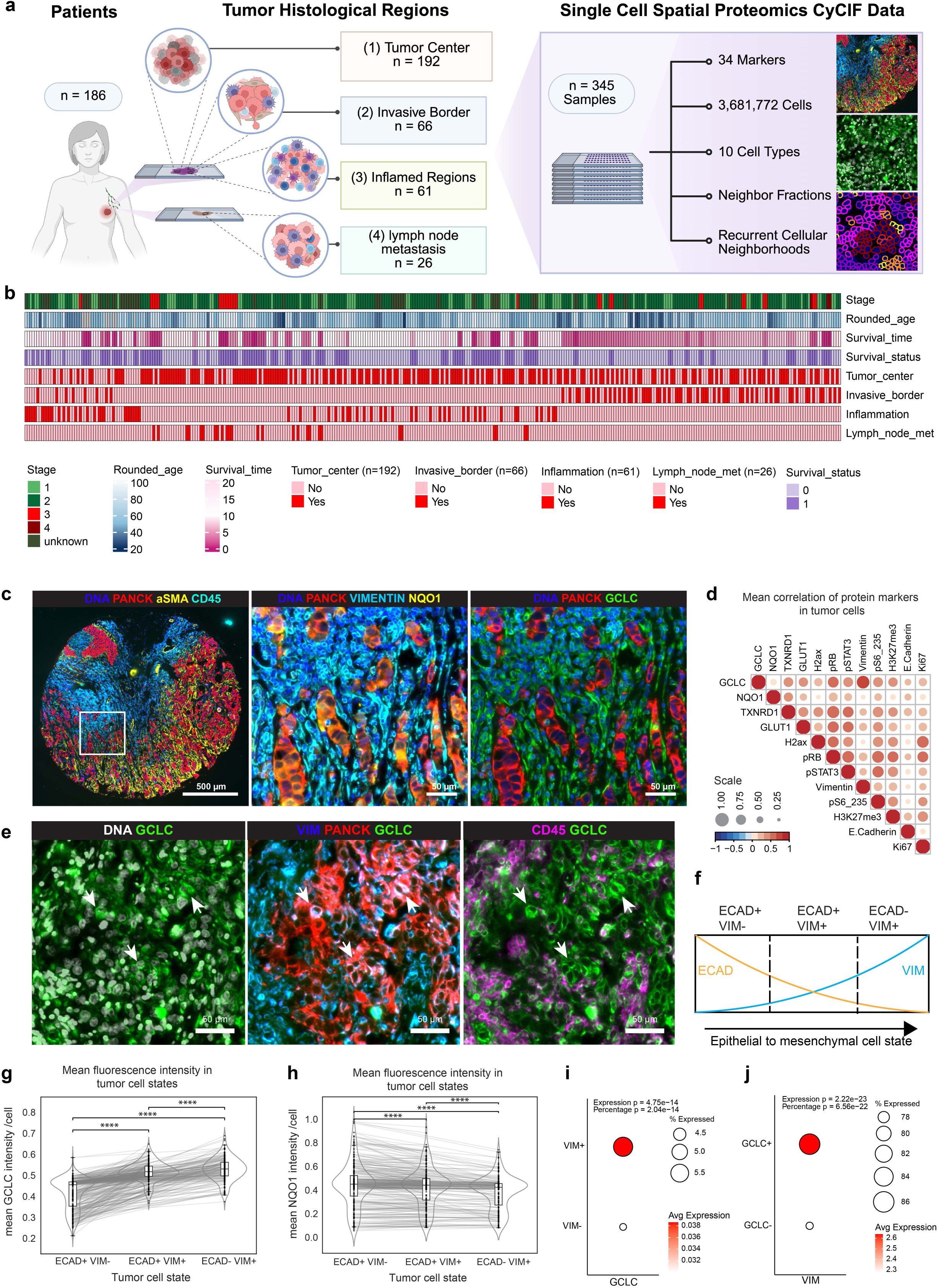
Identification of a tumor cell population characterized by expression of GCLC and VIM. **a** Schematic outlining TMA construction. Different histological regions (1-4) were identified within a tumor, and each histological region was individually cored and selected to be a single sample (n=345) in the TMA. **b** Overview of the primary breast cancer cohort used for spatial profiling and the associated clinical (patient age, disease stage, survival data) and histological characteristics. **c**. A representative TMA core stained for DNA, panCK, aSMA and CD45 (left). The white box illustrates a magnified region stained for NQO1 (middle, yellow) and GCLC (right, green) within tumor (panCK, red) and stromal (VIM, cyan) populations. Scale bars, 500 μm and 50 μm. **d** A Spearman correlation plot of protein expression across tumor cells is presented. All correlations were significant. The dot size and color show the strength and direction of correlation (blue means negative, red means positive). **e** A representative area of GCLC (green) expression in VIM+ (blue) tumor cells (panCK, red) is presented. The left image shows GCLC counterstained with Hoechst (gray), the middle image shows GCLC, VIM, and panCK, and the right image shows GCLC with CD45 (magenta). The overlay of red, green and blue in the middle images appear as white. Likewise, the overlay of green and magenta appear white in the right images. Arrowheads show examples of GCLC+ VIM+ tumor cells. **f** Schematic showing the shift of tumor cells from a more epithelial to a more mesenchymal phenotype by CyCIF marker expression. **g, h** The staining intensity of GCLC (h) or NQO1 (i) in ECAD+ VIM-, ECAD+ VIM+, and ECAD+ VIM+ cells per patient (individual dots) is presented and p-values (paired two-sided t-test) for the indicated comparisons are shown. **i** GCLC expression in VIM+ and VIM-tumor cells from scRNASeq^19^. **j** VIM expression in GCLC+ and GCLC-tumor cells from scRNASeq^19^.

Quantification of NQO1, GCLC, and TXNRD1 expression across the identified cell types revealed that NQO1 was largely expressed in tumor cells. In contrast, GCLC was more commonly expressed in immune and stromal cells, although in some samples over half of tumor cells were GCLC+ (Supplementary Fig. 2e). The proportion of TXNRD1+ cells was low or absent across these cell types (Supplementary Fig. 2e-f). Visual inspection supported enrichment of NQO1 in tumor cells and GCLC in non-tumor cell populations (Fig. 2c). GCLC and TXNRD1 expression in tumor cells decreased with increasing disease stage while NQO1 levels did not (Supplementary Fig. 2g). This observation agrees with earlier studies showing that tumor cells depend on different OSR proteins over the course of disease progression^25^ and suggests that GCLC and TXNRD1 may be important earlier in the disease. Further, consistent with a previous study^26^, we noted a significantly shorter overall survival in patients with high NQO1 expression in tumor cells (Supplementary Fig. 2h).

To reveal associations between OSR proteins and specific cell states, we performed a pairwise correlation analysis for cell state markers across tumor cells. GCLC, NQO1, and TXNRD1 did not correlate strongly with each other (Fig. 2d), in line with the lack of a coordinated NRF2 program observed in the scRNAseq data. The strongest correlation we observed among all of the markers was between GCLC and VIM, suggesting that GCLC expression could be associated with a mesenchymal program. Visual inspection confirmed the presence of GCLC+VIM+ tumor cells in the tumor samples (Fig. 2e). The epithelial-to-mesenchymal transition (EMT) – a process observed in some primary breast tumors^27^ through which epithelial cells begin to downregulate epithelial genes (e.g., *CDH1* encoding E-cadherin, or ECAD) and upregulate mesenchymal markers (e.g., *VIM*) – refers to a spectrum of cell states^28^. Previous reports using mouse and tumor cell line models suggested a reliance of mesenchymal tumor cells on glutathione metabolism and noted elevated GCLC expression levels^29,30^. Moreover, the expression of GCLC was confined to VIM+ tumor cells regardless of the immediate immune and stromal cell neighbors (Supplementary Fig. 2i).

We explored the possibility that GCLC expression in TNBC tumor cells may fall along this spectrum of epithelial-mesenchymal cell phenotypes (Fig. 2f) by comparing GCLC expression across tumor cell populations that were ECAD+VIM-(epithelial), ECAD+VIM+ (intermediate), or ECAD-VIM+ (mesenchymal) and found that GCLC, but not NQO1, is upregulated concurrently with the epithelial-to-mesenchymal phenotypic shift (Fig. 2g-h). Consistently, scRNASeq data revealed that GCLC+ tumor cells had a significantly higher expression of VIM than GCLC-tumor cells (Fig. 2i), with over 85% of GCLC+ tumor cells expressing VIM. Similarly, VIM+ tumor cells had significantly higher expression of GCLC than VIM-tumor cells, though only 5% of VIM+ cells express GCLC (Fig. 2j). Collectively, these results indicate that a population of TNBC tumor cells concomitantly express GCLC and VIM. This is, to our knowledge, the first evidence of GCLC expression in cells with mesenchymal-like phenotypes occurring in patient samples.

In summary, we found that the OSR proteins NQO1 and GCLC are expressed across multiple cell types within the TME, with NQO1 more common in tumor cells and GCLC more common in the stroma and immune populations. These expression patterns varied across tumor cell states, indicating that phenotypic shifts in tumor cells can influence the expression of OSR proteins as is the case with GCLC in VIM+ tumor cell populations.

### GCLC+VIM+ tumor cells reside in inflamed tumor areas, in close proximity to immune cells

We next sought to determine the spatial distribution of OSR proteins across the histological regions. We found that GCLC, NQO1, and TXNRD1 were all highly expressed in tumor cells in inflamed regions (Fig. 3a), and that expression of NQO1 and GCLC was significantly higher in those regions compared to tumor cells within tumor centers (Supplementary Fig 3a).

**Figure 3.**
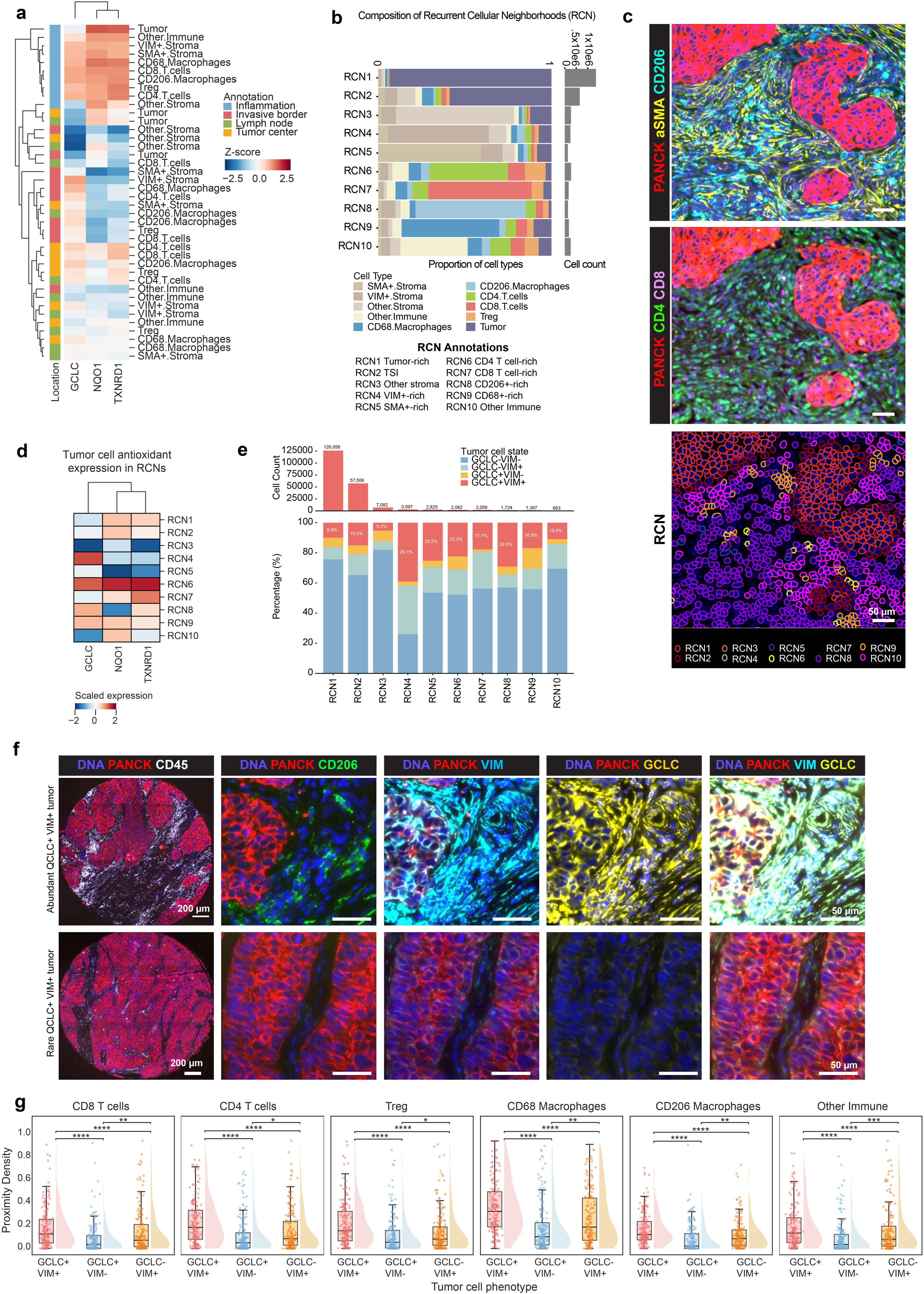
OSR protein distribution in histological regions and recurrent cellular neighborhoods. **a** Heatmap showing the relative protein expression of GCLC, NQO1, and TXNRD1 in different cell types across histological regions. **b** The stacked bar plot shows the proportion of cell types of all cells within recurrent cellular neighborhoods (RCNs). The grey bar plot (right) shows absolute cell counts for each RCN. **c** CyCIF images (top and middle) of a representative area showing cell type presence (panCK = red, SMA = yellow, CD206 = cyan, CD4 = green, CD8 = purple) together with corresponding RCN cell residence (bottom). Scale bars, 50 μm. **d** A heatmap shows the relative protein expression of GCLC, NQO1, and TXNRD1 in tumor cells across the ten RCNs. **e** The stacked bar plot shows the proportion of tumor cells based on their GCLC and VIM expression status. The bar plot above shows the counts of GCLC+VIM+ cells in each RCN. **f** Representative images of GCLC+VIM+ high (top) and low (bottom) cores: a lower magnification image showing complete cores with tumor cells labeled with panCK (red), immune cells with CD45 (white) and cells are counterstained with Hoechst (blue) to show tumor architecture. Scale bar = 200 μm. A higher magnification area (scale bar = 50 μm) from each is presented to show CD206+ myeloid cells (green), VIM (cyan), and GCLC (yellow). **g** Proximity analysis was performed on GCLC+VIM+, GCLC+VIM-, and GCLC-VIM+ tumor cells relative to different immune cell populations. The results are shown as combined bar and violin plots, with the box representing IQR from Q1 to Q3. The horizontal line of the box shows the median, whiskers extend to 1.5 × IQR, and points beyond are plotted as outliers. Individual points represent individual patients. P-values are from Wilcoxon signed-rank test.

To test whether tumor cells expressing a given individual OSR protein are spatially correlated or randomly distributed, we calculated the Moran’s I value for GCLC, NQO1, TXNRD1, and Ki67. This analysis revealed that the OSR protein-expressing tumor cells had a significantly higher degree of spatial autocorrelation (Moran’s I values close to 1, indicative of spatial clustering) when compared to the proliferation marker Ki67 (Moran’s I values approaching 0 indicative of random spatial distribution), regardless of the histological region (Supplementary Fig. 3b). Thus, OSR protein-expressing tumor cells exhibit significant spatial colocalization but distinct expression across histological regions.

To assess whether the presence of OSR protein-expressing tumor cells in inflamed histological regions translates to OSR protein expression in tumor cells in close proximity to immune cells, we identified ten recurrent cellular neighborhoods (RCNs) by clustering the cell types based on the enrichment of neighbors within a radius of 30 microns (see Methods; Fig. 3b, c, Supplementary Fig. 3c). These RCNs captured the major tissue structures such as tumor-rich zones (RCN1), tumor-stroma interface (RCN2), T cell-rich zones (RCN6 CD4+ T cells, RCN7 CD8+ T cells), and myeloid-rich zones (RCN8 CD206+, RCN9 CD68+). Other immune cells were enriched in RCN10. Different stromal cell populations were enriched in RCN4 (VIM+), RCN5 (SMA+), and RCN3 (Other stroma).

Calculating relative OSR protein expression in either all cells (Supplementary Fig. 3d) or tumor cells (Fig. 3d) across the RCNs revealed that all three OSR proteins were highly expressed in T cell rich RCNs 6 and were lowest in RCN3, which is enriched for Other Stroma. NQO1 was most highly expressed in tumor cells from many RCNs, including RCN1, RCN2, RCN6, and RCN9 (Fig. 3d). In contrast, GCLC expression in tumor cells was highest in RCN4 and RCN8, neighborhoods enriched for VIM+ stroma and CD206+ myeloid cells, respectively. These RCNs were also enriched for GCLC+VIM+ tumor cells (Fig. 3e).

A visual inspection of histological regions with a large proportion of GCLC+VIM+ tumor cells confirmed abundant immune cell infiltration near GCLC+VIM+ cells (Fig. 3f). This observation prompted us to calculate the proximity between immune cells and GCLC+VIM+ tumor cells and compare them with the proximity of immune cells to GCLC-VIM+ and GCLC+VIM-tumor cells. We found that GCLC+VIM+ tumor cells were significantly closer to all immune cell types than the other tumor cell populations queried (Fig. 3g). Taken together, our analyses show that certain OSR proteins are enriched in specific cellular neighborhoods. Specifically, NQO1+ tumor cells are distributed broadly across cellular neighborhoods while GCLC+VIM+ tumor cells are frequently near immune cells.

### GCLC+VIM+ tumor cells are characterized by a proliferative, less stressed transcriptional phenotype

Having identified and characterized the spatial distribution of double-positive GCLC+VIM+ tumor cells, we next sought to reveal their gene expression programs using the scRNASeq data. Pathway enrichment analysis revealed that GCLC+VIM+ cells relative to GCLC+VIM-cells exhibited significantly upregulated Epithelial-Mesenchymal Transition (EMT) together with the Hedgehog and TGFB signaling pathways (Fig. 4a; Supplementary Table 6), both of which can promote EMT^31,32^. Inflammatory signaling pathways were also elevated in the GCLC+VIM+ tumor cell population relative to GCLC+VIM-tumor cells. However, tumor cells negative for GCLC but positive for VIM (GCLC-VIM+) had significantly higher expression of inflammatory pathways than GCLC+VIM+ tumor cells indicating a spectrum of inflammatory pathway gene expression (Fig. 4b; Supplementary Table 6). We also observed that the double-positive GCLC+VIM+ cells had high scores for proliferation (Fig. 4c-d), and low gene signature scores for anastasis (Supplementary Table7), a pro-survival pathway^33^ that has been implicated in drug resistance^34,35^. (Fig. 4e-f). These results suggest that GCLC+VIM+ cells are shifted towards EMT, but they are still proliferative and likely not primed for drug resistance^36,37^.

**Figure 4.**
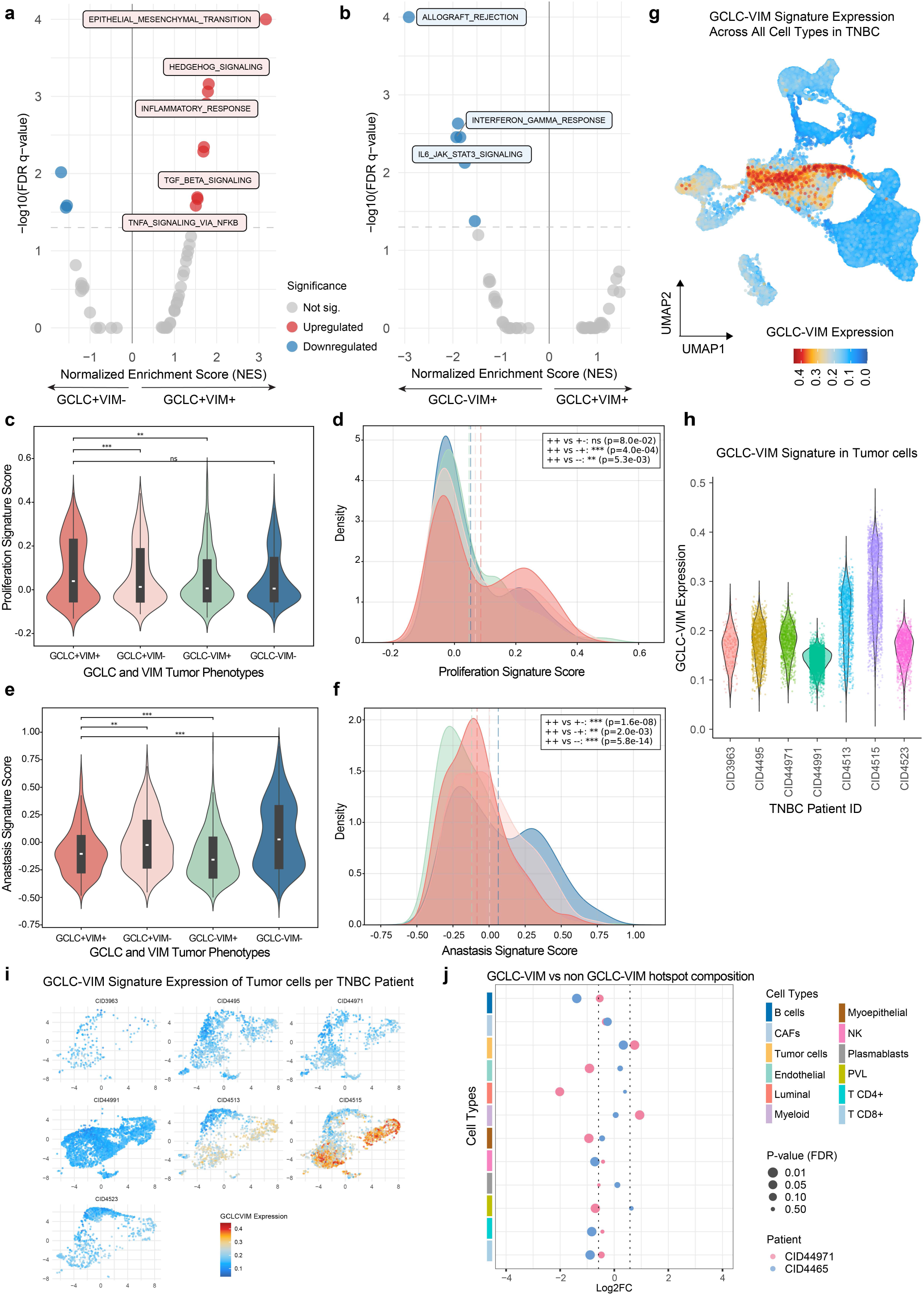
Characterization of the GCLC+VIM+ tumor cell population. **a** A volcano plot shows significantly upregulated (right side) and downregulated (left side) Hallmark pathways in GCLC+VIM+ tumor cells compared to GCLC+VIM-tumor cells. **b** A volcano plot shows significantly upregulated (right side) and downregulated (left side) Hallmark pathways in GCLC+VIM+ tumor cells compared to GCLC-VIM+ tumor cells. **c** Violin plots showing Proliferation signature scores in GCLC+VIM+, GCLC+VIM-, GCLC-VIM+, and GCLC-VIM-tumor cells and statistical significance between indicated comparisons is shown (**p<0.01, ***p<0.001). **d** A density plot showing PCNA signature expression in GCLC+VIM+, GCLC+VIM-, GCLC-VIM+, and GCLC-VIM-tumor cells. **e** Violin plots show anastasis signature expression in GCLC+VIM+, GCLC+VIM-, GCLC-VIM+, and GCLC-VIM-tumor cells and statistical significance between indicated comparisons is shown (**p<0.01, ***p<0.001). **f** A density plot showing anastasis signature expression in GCLC+VIM+ (++), GCLC+VIM-(+-), GCLC-VIM+ (-+), and GCLC-VIM-(--) tumor cells. P-values (**c-f**) are calculated using the Wilcoxon rank-sum test. **g** A UMAP shows all cells from TNBC samples and the expression of the GCLC-VIM signature is overlaid. **h** Violin plots show GCLC-VIM signature expression in tumor cells across TNBC patients. **i** Expression of the GCLC-VIM signature is shown in UMAPs of tumor cells by individual patients. **j** The cell type composition of GCLC-VIM signature hotspots and non-hotspots was compared using log fold change and plotted for each slide.

To assess how the expression of GCLC and VIM relates to the AntiOx signature, we developed a tumor cell-specific, 104-gene “GCLC-VIM” signature based on highly expressed genes in GCLC+VIM+ tumor cells (Methods, Fig. 4g, Supplementary Table 8). Interestingly, the GCLC-VIM signature score varied across TNBC patients unlike what was observed with the AntiOx signature. Whereas all seven patients had comparable AntiOx signature scores, the distribution of GCLC-VIM signature scores in patients CID4513 and CID4515 trended higher compared to the other patients (Fig. 4h). As with the AntiOx signature, the GCLC-VIM signature was expressed in multiple tumor cell clusters (Fig. 4i). We observed a significant, although modest correlation between the AntiOx and GCLC-VIM signatures across tumor cells in the scRNASeq data, indicating that the GCLC-VIM signature is only partially associated with OSR gene expression (Supplementary Fig. 4a) further highlighting the distinct characteristics of GCLC-VIM tumor cells.

Next, we identified GCLC-VIM signature hotspots in the spatial transcriptomics data, noting that GCLC-VIM signature hotspots exhibited distinct regions of signature expression compared to AntiOx (Supplementary Fig. 4b-d compared with Fig. 1f, g). Similarly to the AntiOx signature, the GCLC-VIM tumor signature hotspots were dominated by tumor cells (Supplementary Fig. 4c). Myeloid cells, myoepithelial cells, and CAFs were also highly represented in the GCLC-VIM signature hotspots (Supplementary Fig. 4c, d). When comparing GCLC-VIM signature hotspots and other spots composition across patients, we found that only tumor cells were consistently increased in the GCLC-VIM signature hotspots (Fig. 4j). Collectively, these results suggest that the GCLC+VIM+ cells do not simply represent a mesenchymal subset of Antiox high cells but rather are a phenotypically and spatially distinct tumor cell population.

### GCLC+VIM+ tumor cells are associated with better patient outcomes

The lower expression of anastasis genes and an increased proliferation signature in GCLC+VIM+ tumor cells suggest these cells may be more susceptible to chemotherapies. As a functional test of GCLC+VIM+ tumor cell sensitivity to first line therapies, we assessed how the mesenchymal state and GCLC expression relate to drug sensitivity in TNBC cell lines from the LINCS dataset^38^. We used the Basal A and Basal B subtypes^39^ as surrogates for the mesenchymal, VIM+ cell state, with Basal B classified as a more mesenchymal state (Fig. 5a). Basal B cells treated with cisplatin showed significant resistance based on the GRMax, though not other metrics (Fig. 5b), suggesting that the molecular subtype does not strongly affect drug sensitivity. To determine whether elevated GCLC expression in Basal B cell lines enhances chemosensitivity, we correlated GCLC expression to GR metrics. This analysis revealed that GCLC expression significantly associated with cisplatin sensitivity (Fig. 5c). Thus, tumor-cell GCLC expression associated with improved sensitivity to chemotherapy, an observation with potential clinical relevance for patient outcomes.

**Figure 5.**
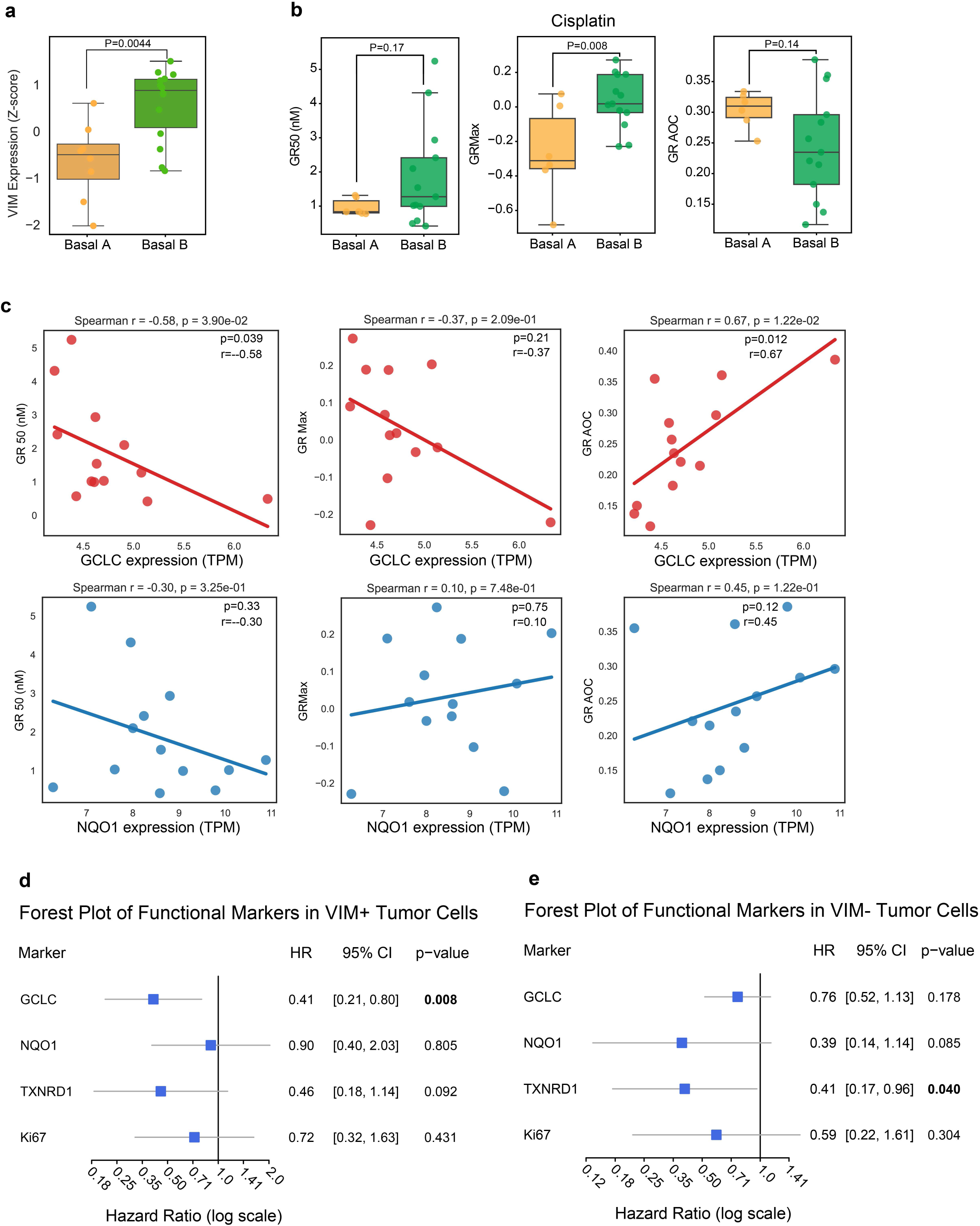
GCLC+VIM+ tumor cells are associated with sensitivity to chemotherapy. **a** Bar plot showing VIM gene expression in Basal A and Basal B TNBC subtypes. **b** Bar plots showing GR50, GRMax, and GRAOC of Basal A and Basal B TNBC subtypes for cisplatin. P-values in a and b were calculated using an unpaired, two-tailed t-test. Horizontal line shows median, box edges show upper and lower quartiles, and whiskers extend to 1.5 × IQR. **c** Correlation plots of cisplatin sensitivity with GCLC (top row) or NQO1 (bottom row) expression in TNBC cell lines. GR_Max_ (left), GR_50_ (middle), and GR_AOC_ (right) metrics are presented. Spearman’s r and p-values are shown on the corresponding graphs. **d** Forest plot showing hazard ratio for OSR protein or Ki67 expression in VIM+ tumor cells in inflamed tumor spatial compartments. **e** Forest plot showing hazard ratio for OSR protein and Ki67 expression in VIM-tumor cells in inflamed tumor spatial compartments.

To test the outcomes of TNBC patients with GCLC+VIM+ tumor cells, we performed Cox regression analysis. Given that GCLC+VIM+ tumor cells were more frequently found in areas of inflammation, we selected inflamed histological regions for this analysis. We found that VIM+ tumor cells in these areas were associated with a longer overall survival if they also expressed GCLC (See Methods; Fig. 5d). GCLC expression in VIM-tumor cells did not reveal any survival benefit (Fig. 5e), suggesting that both VIM and GCLC positivity are required for this phenotypic survival association.

In summary, our work has shown that OSR pathway members are differentially expressed across tumor cell populations and in diverse microenvironments. The expression of some of these proteins, like GCLC, may be primarily expressed by immune and stromal cell types. However, altered tumor cell state, such as a shift to a more mesenchymal state, may also lead to their expression in malignant cells. Furthermore, our results suggest that the expression of OSR genes and proteins in specific malignant cell populations may mark chemo-responsive cells.

## Discussion

Using a large spatial proteomics dataset from 186 TNBC patients combined with re-analysis of public single-cell and spatial transcriptomics datasets, we comprehensively spatially mapped OSR gene and protein expression in the TME of TNBC. We uncovered a tumor cell state defined by GCLC and VIM expression characterized by lower expression of anastasis-related genes and higher expression of proliferation-related genes compared to other tumor cell populations. GCLC+VIM+ cells were also spatially located in close proximity to diverse immune populations. Importantly, the GCLC+ VIM+ tumor cell state predicted sensitivity to chemotherapy in vitro, and was associated with prolonged overall survival in patients with primary TNBC. These analyses provide insights into the spatial distribution and cell type specificity of OSR proteins in TNBC.

Drugs targeting OSR proteins have been studied in clinical phase I trials (NQO1^11^, TXNDR1^12^, GCLC^13,14^). Overall, these drugs are reasonably well-tolerated; however, their efficacy as anti-cancer agents varied. Combining OSR protein-targeting drugs with oxidative stress inducing therapies, such as radiation therapy, could theoretically offer synergistic benefit^40,41^. However, the cell type specificity of OSR proteins has remained largely unexplored. Our study shows that NQO1 is primarily expressed in tumor cells, suggesting NQO1 inhibitors would have a minimal impact on non-tumor cells in the TME. In line with prior reports^26^, we show that high tumor NQO1 expression is associated with poor clinical outcomes. Together, these findings support further exploration of NQO1 inhibition in tumors highly expressing this protein.

In this study, we found high GCLC expression in immune and stromal cell populations suggesting that glutathione may be a key antioxidant in the TME. This might offer a plausible explanation why clinical trials have failed to demonstrate efficacy for GCLC inhibitor buthionine sulfoximine (BSO). Effector T cell responses are particularly shown to be dependent on glutathione^42^, which requires functional GCLC (GCLC being the rate-limiting step in the glutathione synthesis). In myeloid cells, increased ROS from BSO or chemotherapy can elevate PD-L1 expression, promoting an immunosuppressive tumor milieu^43^. Although high GCLC expression in tumor cells is associated with metastasis and therapy resistance^44,45^, GCLC inhibition may weaken immune surveillance more than tumor cells, ultimately favoring tumor persistence. Thus, our work highlights the intricate biology of distinct OSR proteins in TNBC, and suggests that, in contrast to NQO1, systemic targeting of GCLC does not represent a promising therapeutic approach.

Beyond understanding how the distribution of OSR proteins across cell types might affect their efficacy as therapeutic targets, our study provides evidence that chemotherapy sensitivity might be related to OSR expression. Our analysis shows that unlike NQO1, GCLC was expressed in only a subset of tumor cells, most of which were in a mesenchymal-like state, expressing VIM. Although GCLC+VIM+ cells are shifted towards EMT, they are still proliferative, and therefore likely not primed for chemotherapy resistance. Indeed, this tumor cell population was associated with enhanced chemotherapy responses and improved patient outcomes. The finding that high GCLC expression in cell lines correlates with cisplatin sensitivity was surprising, given previous reports [refs]. However, GCLC may be part of a broader cell state that is susceptible to chemotherapy rather than being the driver of drug response. Mechanistic studies that systematically knock out or overexpress GCLC could establish whether this relationship is causal, but are outside the scope of this study. Our spatial analysis showed close proximity of GCLC+VIM+ tumor cells to diverse immune and stromal cell populations, which could suggest that this state may be induced by neighboring cell types. Cancer-associated fibroblasts (CAFs)^46,47^, CD8 T cells^48,49^ as well as different myeloid cell populations^50–52^ can induce EMT through secreted factors in breast as well as other cancer types. However, NQO1+ tumor cells, despite being found in several immune-rich RCNs, did not show a tendency towards a mesenchymal phenotype. The spatial hotspots and niches likely also have significant consequences for the maintenance of distinct tumor cell populations. For example, GCLC+VIM+ tumor cells were spatially associated with CD8+ T cells, which may exert cytotoxic functions. The GCLC+VIM+ cells were also proximal to CD206+ macrophages, which in the breast are tissue-resident macrophages not associated with inflammation^53,54^ and rather infiltrate less dense breast tissue, where they potentially modulate the surrounding stroma to resemble the normal, non-cancerous stroma^55^. These observations point to the possibility that GCLC+VIM+ tumor cells reside in microenvironments that are less conducive to tumor cell expansion.

While our study provided insights into the spatial distribution of OSR proteins in TNBC, there are several limitations of our study that could be expanded on in future studies. Firstly, we used a collection of tumor samples using a TMA, an approach that enabled us to explore OSR protein expression at a much larger scale than would be feasible with whole-tissue spatial profiling^56^. To partially account for the spatial heterogeneity, these TMAs were sampled from different locations within each tumor, allowing evaluation of histologically distinct tumor regions and endeavoring to capture the spatial heterogeneity of TNBC tumors. Secondly, the antibody panel for CyCIF was limited to the selected immune populations and tumor phenotypic markers, and spatial transcriptomics used in our work was not of single-cell resolution. Finally, the role of GCLC in patient prognosis needs to be interpreted with caution, since similar large, annotated single-cell spatial patient cohorts were unavailable for external validation, in particular with populations of known increased incidence of TNBC, like BRCA1 mutation carriers and African American women.

In conclusion, our results demonstrate that the spatial distribution of OSR genes and proteins in tumors is diverse and can have important clinical implications. The data from this large, well-annotated cohort, provide a foundation for future studies to explore OSR protein expression at single-cell and spatial resolution. These approaches will pave the way to a deeper understanding and therapeutic exploitation of OSR pathways in cancer.

## Methods

### Transcriptomics

#### Reference to scRNASeq dataset used

We used a subset of a published scRNAseq breast cancer dataset^19^, which included 11 ER⁺ and 10 TNBC patients, to study patterns of OSR gene expression in tumor cells. After initial exploration and quality control, we excluded patients with fewer than 150 cancer epithelial cells. The final dataset comprised 66,076 cells from 16 patients, including 9 ER⁺ and 7 TNBC cases. Data normalization and rescaling were performed using the R package Seurat v5.2.1^57^, and visualization was conducted using UMAP.

#### AntiOx signature derivation

The AntiOx signature was generated from a curated set of 86 genes with established OSR functions^58^. Single-cell module scores were calculated using the ‘*AddModuleScore_UCell’* function from the *UCell* package^59^ with default parameters.

#### Gene Set Enrichment Analysis

To characterize pathways across tumor subclusters, we performed GO-based Gene Set Enrichment Analysis (GSEA) using gseGO^60^. For each of the six malignant clusters, we selected the top 5 enriched GO pathways and then plotted their Normalized Enrichment Scores (NES) across all six clusters to enable cross-cluster comparison. To reduce redundancy and assign concise biological themes per cluster, we computed pairwise semantic similarity among enriched terms using pairwise_termsim and summarized them with treeplot^61^, from which we derived a compact set of representative terms for each cluster.

To identify pathways associated with oxidative stress, GSEA was performed based on DEGs between high and low oxidative stress groups (with a threshold of oxidative stress scores > 0.15). Differential expression analysis was conducted using the ‘*FindMarkers’* function of Seurat with the Wilcoxon rank-sum test to compare two groups. Genes with an adjusted p-value < 0.01 and absolute values of log₂ fold change > 0.5 were considered significantly differentially expressed. Downstream analysis was performed using the *gseapy* package (version 1.1.8)^62^ with gene sets provided in GMT format (h.all.v2024.1.Hs.symbols.gmt) in Python. permutation_num was set as 1,000 to estimate significance and a random seed as 42 to ensure reproducibility. Normalized Enrichment Scores (NES) were computed for each gene set, pathways with FDR < 0.05 and positive NES were classified as upregulated, while those with FDR < 0.05 and negative NES were classified as downregulated. Non-significant pathways were labeled accordingly. GSEA results with AntiOx^High^ vs AntiOx^Low^ are in Supplementary Table 3 and different GCLC-VIM group comparisons are in Supplementary Table 6.

#### Spatial transcriptomics

We profiled 2,352 spots from two TNBC Visium slides (slide 1:CID44971; slide 2: CID4465), paired with scRNAseq data from Wu et al^19^. Cellular deconvolution was performed using Cell2location with the full scRNAseq dataset as a reference^24^. Preprocessing and normalization were conducted using the *ScanPy* package^63^. To reduce granularity, the 41 cell types identified by deconvolution were grouped into 12 broader categories: B cells, CAFs, endothelial cells, luminal cells, myeloid cells, myoepithelial cells, PVL cells, CD4⁺ T cells, CD8⁺ T cells, tumor cells, NK cells, and plasmablasts. Spot-level data were subsequently normalized and rescaled for downstream analysis. We used *SpottedPy*^23^ to compute hotspots for AntiOx and GCLC-VIM-derived signatures previously computed, as well as for each of the 12 major cell types, using default parameters. SpottedPy leverages the Getis-Ord Gi* statistic to identify significant stable spatial clusters of locally elevated biological scores of gene expression. Distances between the AntiOx and GCLC+VIM+ hotspots (used as primary variables) and cell type hotspots (comparison variables) were calculated for each slide individually, using a neighborhood parameter of 10. Statistics and plots were derived using R Computing Environment version 4.4.2 [(R Core Team (2021). R: A language and environment for statistical computing. R Foundation for Statistical Computing, Vienna, Austria. URL https://www.R-project.org/.)]

#### GCLC-VIM signature derivation

We utilized cell module scores to quantify the extent of individual cell expression of GCLC/VIM. For this purpose, we used the ‘*AddModuleScore_UCell’* function from *UCell* with default settings^59^. To define the gene signature for GCLC and VIM expression, we extracted gene expression data from the Seurat object and calculated the median expression levels for each gene across all tumor cells. Cells were then classified into four categories based on their expression relative to the median thresholds: (I) “++” for high expression of both genes; (II) “+-” for high GCLC expression and low VIM expression; (III) “-+” for low GCLC expression and high VIM expression; (IV) “--” for low expression of both genes. After classification, we identified the genes defining each subset by calculating differentially expressed genes (DEGs) through the Wilcoxon rank-sum test through the ‘*FindAllMarkers’* function in Seurat. GCLC+VIM+ gene list is in Supplementary Table 8.

#### Proliferation and anastasis scores

Proliferation and anastasis were computed using the ‘*AddModuleScore’* function of Seurat (version 5.2.1) in R for each cell. Anastasis^64^ and proliferation^65^ gene signatures were defined based on published literature. In order to reduce the impact of differences in number between samples, we randomly selected 500 cells from each group for visualization analysis. Proliferation and anastasis signatures are in Supplementary Table 7.

### Cyclic immunofluorescence and image processing

#### AURIA TMA Construction

The TMA was obtained from the AURIA Biobank, Turku, Finland (Decision AB19-1654). This TMA contains over 180 TNBCs collected from a single Organization of European Cancer Institutes (OECI) accredited cancer center with more than ten years of follow up data, including information on tumor stage, patient age, and over ten years of longitudinal data on outcome. The cores were 1.5 mm enabling a large sample from the tumor spatial locations. Furthermore, multiple cores were taken from a single patient in different sample locations enabling intra-tumor comparisons to understand phenotypic differences of the tumor center, invasive border, inflamed regions, and lymph node metastases. Patient metadata can be found in Supplementary Table 4.

#### CyCIF Staining

Tissue-based Cyclic immunofluorescence (t-CyCIF) was performed as previously described^66^, following the protocol available on protocols.io (https://doi.org/10.17504/protocols.io.bjiukkew). Formalin-fixed, paraffin-embedded (FFPE) tissue sections were processed using the BOND RX Automated IHC/ISH Stainer (Leica Biosystems). Slides were baked at 60°C for 30 minutes, dewaxed with Bond Dewax solution at 72°C, and subjected to antigen retrieval using Epitope Retrieval 1 (Leica) at 99°C for 20 minutes. Tissue sections underwent iterative cycles of antibody staining, imaging, and fluorophore inactivation. Antibodies were incubated overnight at 4°C in the dark/humid box (panel in Supplementary Table5). The nuclear counterstaining was performed using Hoechst 33342 (ThermoFisher) mixed with indicated antibodies (1ug/ml final concentration). Slides were then wet-mounted with 100 μL of 10% glycerol in PBS and imaged using a CyteFinder slide-scanning fluorescence microscope (RareCyte Inc., Seattle, WA) equipped with a 20× objective lens (0fw.75 NA). Fluorophore inactivation between cycles was achieved by incubating the slides in a solution of 4.5% hydrogen peroxide and 24 mM NaOH in PBS under LED illumination for 1 hour.

#### Image Processing and Quality Control

Image analysis was performed using the MCMICRO^67^ pipeline and customized scripts in Python (available in GitHub: https://github.com/labsyspharm/mcmicro). Raw images were stitched and registered using the ASHLAR^68^ module of MCMICRO.The assembled OME.TIFF files from each slide were segmented using that params.yaml file in the Github repository and quantified using UNMICST2^69^ to generate single-cell data. More details and source code can be found at https://www.cycif.org and as listed in the software availability section. Quality of the imaging data was assessed visually. All slides were reviewed for tissue integrity and accurate registration. Any areas with artifacts, deformed or lost tissues, poor registration, or cores with fewer than 100 cells were excluded from downstream analysis. Segmentation was iteratively checked and adjusted to improve accuracy based on segmentation masks. Antibodies that did not give the expected staining pattern or had very low signal were excluded from analysis. Marker gating was performed manually with visual inspection for each marker using Gater (https://github.com/labsyspharm/gater). Marker intensities were then normalized using the identified gates such that a value above 0.5 was considered positive.

#### Cell type calling

Celltype calling was performed using Scimap^70^ in Python. We used *’sm.pl.gate_finder’* to find the gate values for each marker per image, followed by using *’sm.pp.rescale’* to standardize all marker expression to a common scale between 0 and 1. Here values above 0.5 will be considered positive, otherwise, it will be considered as a negative value. Cell type calling was performed iteratively using the function *‘<u>sm.hl</u>.classify’*, first dividing tumor (panCK+ or E.Cadherin+), macrophage (CD68+ or CD206+), other immune cells (CD45+), and myofibroblasts (SMA+), followed by the division of macrophages into subpopulations: CD68+ macrophages (CD68+) and CD206+ macrophages (CD206+), and other immune cells into subpopulations: CD8+T-cells (CD8a+), CD4+T-cells (CD4+) and FOXP3+CD4+T-regs (FOXP3+ and CD4+) as well as other immune cells (CD45+ but negative for the other immune cell markers, possibly including B-cells, NK-cells, dendritic cells and granulocytes). Cells labeled as “other” were negative for the aforementioned markers. In our dataset, CD45 was not strongly expressed by macrophages, which is why macrophages were classified separate from the other immune cells. In addition, a subset of macrophages expressed CD4, which is why macrophages were classified prior to T-cells. The resulting cell type calls were inspected on top of the images using the function *‘<u>sm.pl</u>.image_viewer*’ and parameters were adjusted accordingly.

### Image analysis

#### Protein correlations in CycIF data

Spearman rank correlation analysis was used to evaluate pairwise associations between proteins, including ‘Ki67’, ‘GCLC’, ‘NQO1’, ‘TXNRD1’, ‘GLUT1’, ‘H2ax’, ‘pRB’, ‘pSTAT3’, ‘Vimentin’, ‘pS6_235’, ‘H3K27me3’, ‘E.Cadherin’. The correlation values and p-values were calculated using the *‘rcorr’* function in the Hmisc package (version 5.2.3) of the R language.

#### Violin plots of marker expression

Following the classification of different tumor cells using a threshold of 0.5, VIM+ECAD+, VIM+ECAD-, and VIM-ECAD+ tumor groups were selected. GCLC and NQO1 expressions were compared with different groups for each patient.

#### Survival Analysis

Survival analyses were performed using Kaplan-Meier plots and associated log-rank p-values (p<0.05) based on NQO1+ tumor cells proportion per patient. NQO1+ tumor cells were defined as those with expression levels above the threshold of 0.5. Patients were stratified into top 25% and bottom 25% quartiles based on the proportion of NQO1+ tumor cells. The plots were made for each filtered annotation separately: inflammation, tumor center and invasive border. Analyses were done using R (version 2025.05.1+513) with the survival package. Survival status and time were included as variables. Plots were visualized with confidence intervals and risk tables with the survminer package.

#### RCN identification

Recurrent cellular neighborhoods (RCNs) were performed using Scimap^70^ in Python. ‘*sm.tl.spatial_count’* was used to compute a neighborhood matrix for each cell centroid with a radius=45px (1 px=0.65 microns), accompanied by clustering cells with similar spatial features through ‘*sm.tl.spatial_cluster’* with k=10 to form RCNs. Selection of the number of clusters was done with the Elbow method (Supplementary Fig. 3c).

#### Proximity Density

The spatial proximity of tumor cells expressing different GCLC-VIM types and different immune cells was performed using ‘*sm.tl.spatial_pscore’* of the Scimap^70^ in Python. Proximity density was selected here, which reflects the ratio of identified interactions to the total number of cells of the interacting types, providing insight into how frequently these cell types interact relative to their population size. The score for each tumor-immune pair was calculated per core with a radius of 30 microns, followed by the average to get the value for each patient. Both the thresholds of GCLC+ and VIM+ were 0.5.

#### Moran’s I

Moran’s I scores were calculated using the *gr.spatial_autocorr* function in the Squidpy Python package per each sample^71^. Visualization was done using S*eaborn* and *Matplotlib* packages in Python, displayed as violin plots.

#### Giotto

Spatial enrichment of cell-cell interactions between defined immune cells and tumor subpopulations were performed using the Giotto R package^72^. For each sample, a Delaunay triangulation was constructed to define cell–cell proximity networks based on centroid positions with a radius of 30 microns, and fold change values per tumor cell type for GCLC expression based on the spatial interaction vs non-interaction with another cell type were calculated using ‘*FindInteractionChangedFeats’* function. The resulting enrichment scores were aggregated across tissue regions (tumor center, invasive border, inflammation, lymph node met).

#### Cox proportional hazards regression model

Tumor cells were defined as VIM+ and VIM-with the threshold of 0.5 of the expression. The median expression of four functional markers (GCLC, NQO1, TXNRD1, Ki67) were computed across different sample locations and types.

Marker expression levels were stratified into quintiles using the ‘*ntile’* function with tile_number=5. Cox proportional hazards regression models were fitted for each marker using the *’coxph’* function, with marker expression (quintile), stage, and a time-dependent age term (including cubic age transformations) as covariates. Patients with unknown stage were excluded. Results were visualized using forest plots with hazard ratio (HR), 95% confidence intervals (CI), and p-value, highlighting statistically significant markers.

### Cell line analysis

Plots pertaining to cell line analyses in Fig. 5 were generated using the SciPy package to calculate the Wilcoxon-rank sums test for the bar graphs, and the Pearson coefficient and p value for the gene expression-signature expression correlations. Visualization was done with the matplotlib.pyplot and seaborn packages. Growth rate inhibition (GR) metrics and cell line metadata were obtained from previously published dose response data^38^, and transcripts per million (TPM) expression data were obtained from the Broad Dep Map portal (DepMap, Broad (2025). DepMap Public 25Q2. Dataset. https://depmap.org^73^). Datasets were filtered to include only results for overlapping triple negative breast cancer cell lines. Linear fits for visualization were generated using the numpy polyfit function, and Spearman correlations were calculated using the spearmanr function in the scipy stats library.

## Code availability

All scripts used for data analysis and to generate the figures have been uploaded to Github: https://github.com/farkkilab/Auria_project/tree/Publication.

## Acknowledgements

We would like to thank Joan S. Brugge for insightful discussions and feedback on the project, and Isaac S. Harris for a critical reading of the manuscript. Jeremy Muhlich provided MCMICRO support. We are grateful for Zoltan Maliga’s antibody validation efforts. This work was supported by the Ludwig Cancer Center at Harvard Medical School, the Gray Foundation, and an ASPIRE Award from the Mark Foundation for Cancer Research. TV is supported by a Research Scholar Grant, PF-24–1316850-01-CD, from the American Cancer Society.

## Notes

https://github.com/farkkilab/Auria_project/tree/Publication.

### Competing Interest Statement

Peter K. Sorger is a co-founder and member of the BOD of Glencoe Software, member of the SAB for RareCyte, Reverb Therapeutics and Montai Health, and consultant for Merck; he holds equity in Glencoe and RareCyte. The other authors declare no potential conflicts of interest.

