## Supplemental Figures for "Integrative single-cell and spatial mapping of oxidative stress response uncovers GCLC+ mesenchymal tumor cell state linked with favorable outcomes in triple negative breast cancer"

**Supplementary Figures**

### Supplementary Figure 1

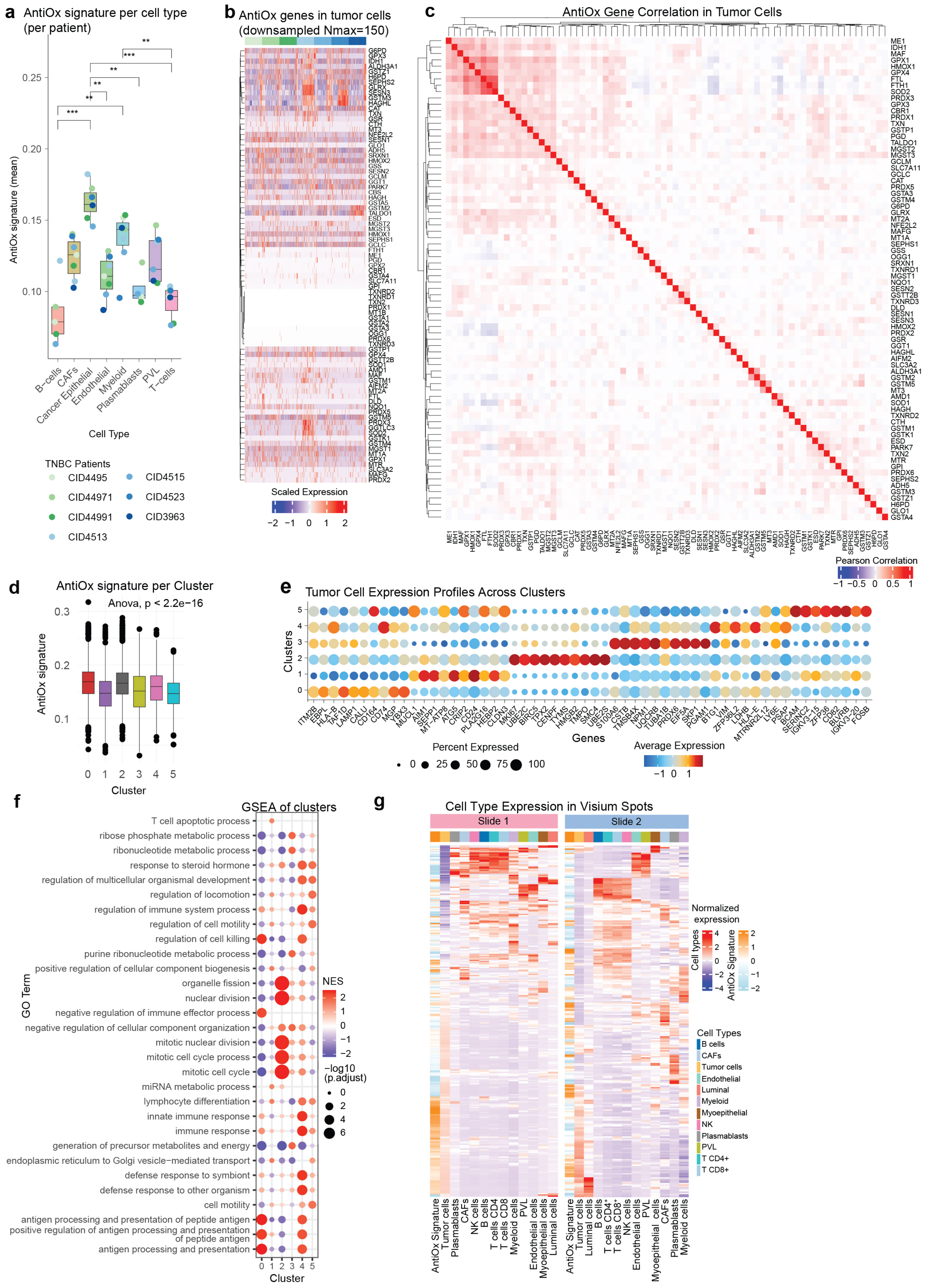

Supplementary Figure 1. **a** Boxplot of the AntiOx signature expression across cell types in TNBC per patient. The box represents the IQR from Q1 to Q3, with the horizontal line indicating the median. Whiskers extend to 1.5 × IQR from the quartiles. Significance values were computed based on Dunn's test after a Kruskal–Wallis test and are shown: ** $p<0.01$, *** $p<0.001$. **b** Heatmap showing OSR gene expression in TNBC tumor cells, clustered by patient. Subsampling of 150 tumor cells per patient was performed. **c** Correlation heatmap showing OSR gene correlation in tumor cells across all TNBC samples. **d** Boxplot ofAntiOx signature score within TNBC tumor subclusters. **e** Dotplot of scaled average gene expression of the top ten most highly expressed genes in the 6 TNBC tumor subclusters. **f** Dot plot showing pathway enrichment of the top 5 significant GO terms for each TNBC tumor subcluster, along with their expression across all clusters. **g** Heatmaps for each Visium slide of normalized AntiOx signature expression and distribution of cell types across the analyzed spots.

Supplementary Figure 2

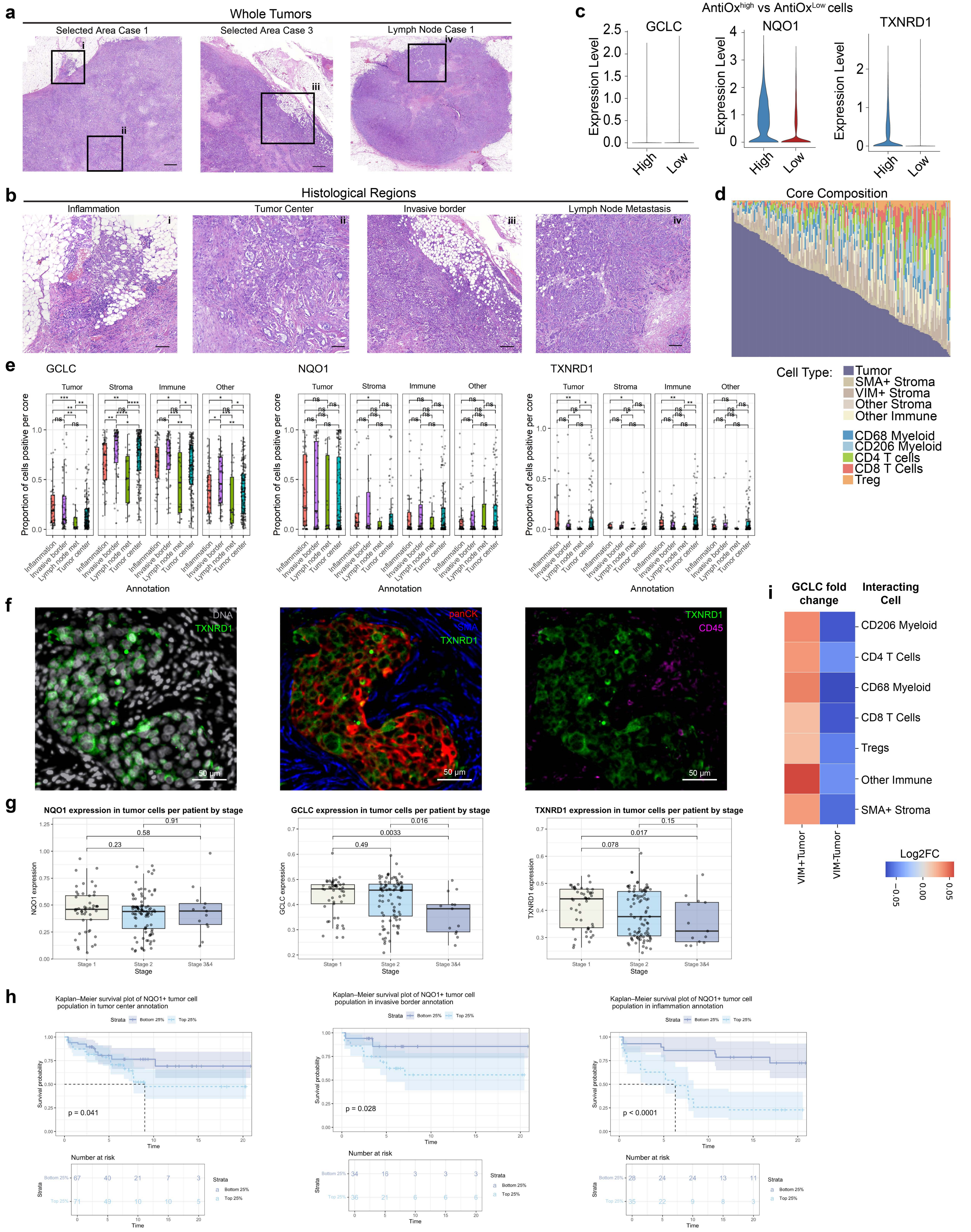

Supplementary Figure 2. **a** Representative H&E showing the annotations for histologic regions (inflammation, tumor center) and the corresponding lymph node metastasis. **b** Violin plots of GCLC, NQO1 and TXNRD1 expression levels in AntiOx high and AntiOx low cells. **c** A waterfall plot of cell type distribution by patients. **d** Quantification of OSR protein staining intensity across histological regions and broad cell types, including tumor, immune, stroma (SMA+), and Other. **e** A representative image of TXNRD1 (green) staining. The left image is counterstained with Hoechst (gray), the middle panel includes panCK (red) and SMA (blue), and the right panel includes CD45 (magenta). **f** OSR protein expression by stage is presented with p-values of indicated comparisons shown. The box represents the IQR from Q1 to Q3, with the horizontal line indicating the median. Whiskers extend to 1.5 × IQR from the quartiles. The dots represent patients and p-values from the Wilcoxon rank-sum test. **g** Patient survival by NQO1 expression in different histological regions (invasive border, top; tumor center, middle; inflammation, bottom). **h** A heatmap showing the log2 fold change in expression of GCLC in either VIM+ or VIM- tumor cells interacting with the indicated immune cells as compared to non-interacting cells. The values were calculated using the Giotto package (see Methods).

### Supplementary Figure 3

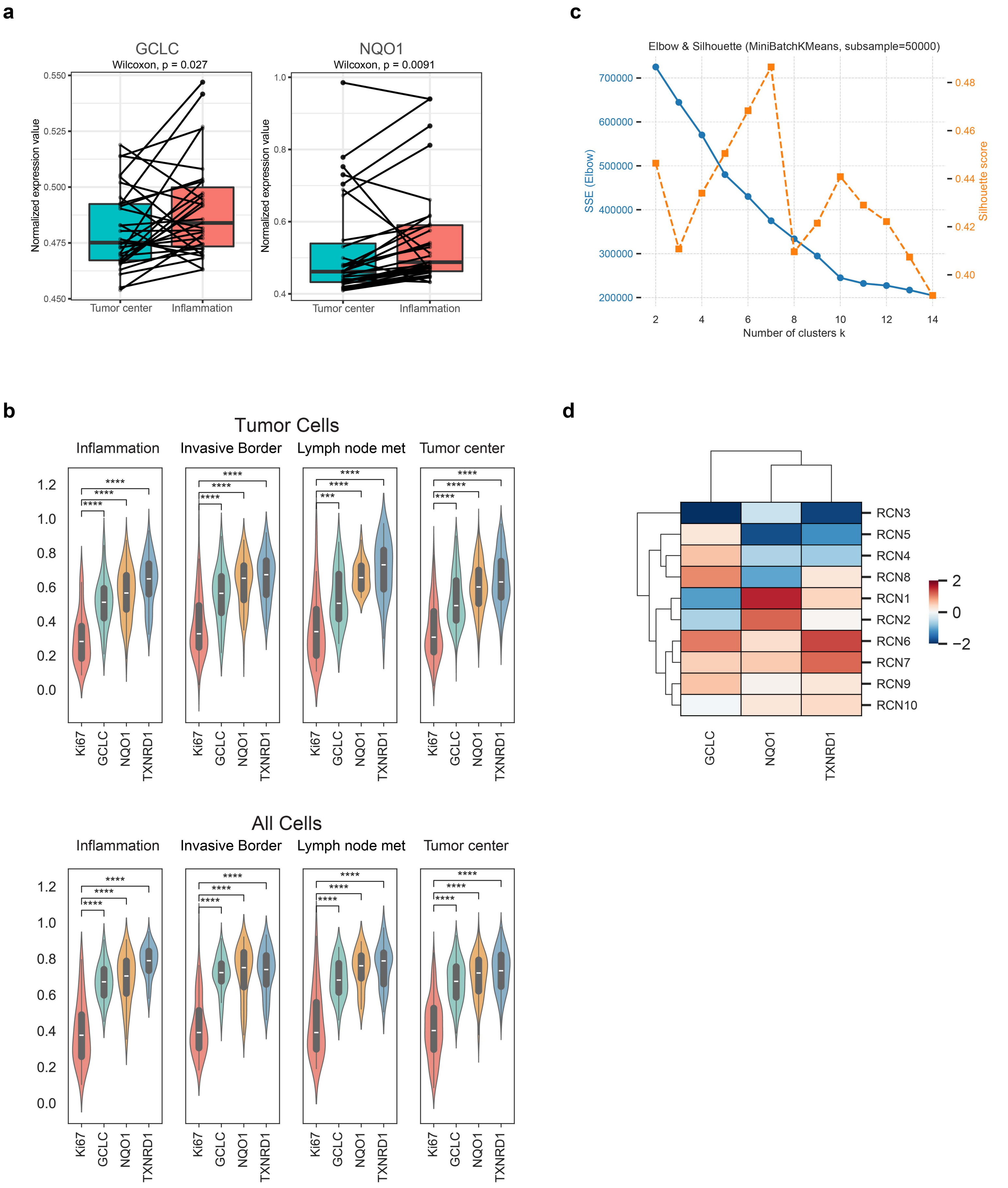

Supplementary Figure 3. **a** Bar plots showing GCLC (left) and NQO1 (right) expression in tumor center or inflamed cores in matched patients. Significance was calculated using Wilcoxon signed-rank test. **b** Moran's I values are plotted for the OSR proteins and Ki67 in tumor cells (top) or all cells (bottom) across tumor spatial compartments. P-values are from Wilcoxon signed-rank test: *** $p<0.001$, **** $p<0.0001$. **c** An elbow plot for determining the optimal number of RCN clusters. **d** A heatmap shows the relative protein expression of GCLC, NQO1, and TXNRD1 in all cells across the ten RCNs.

Supplementary Figure 4

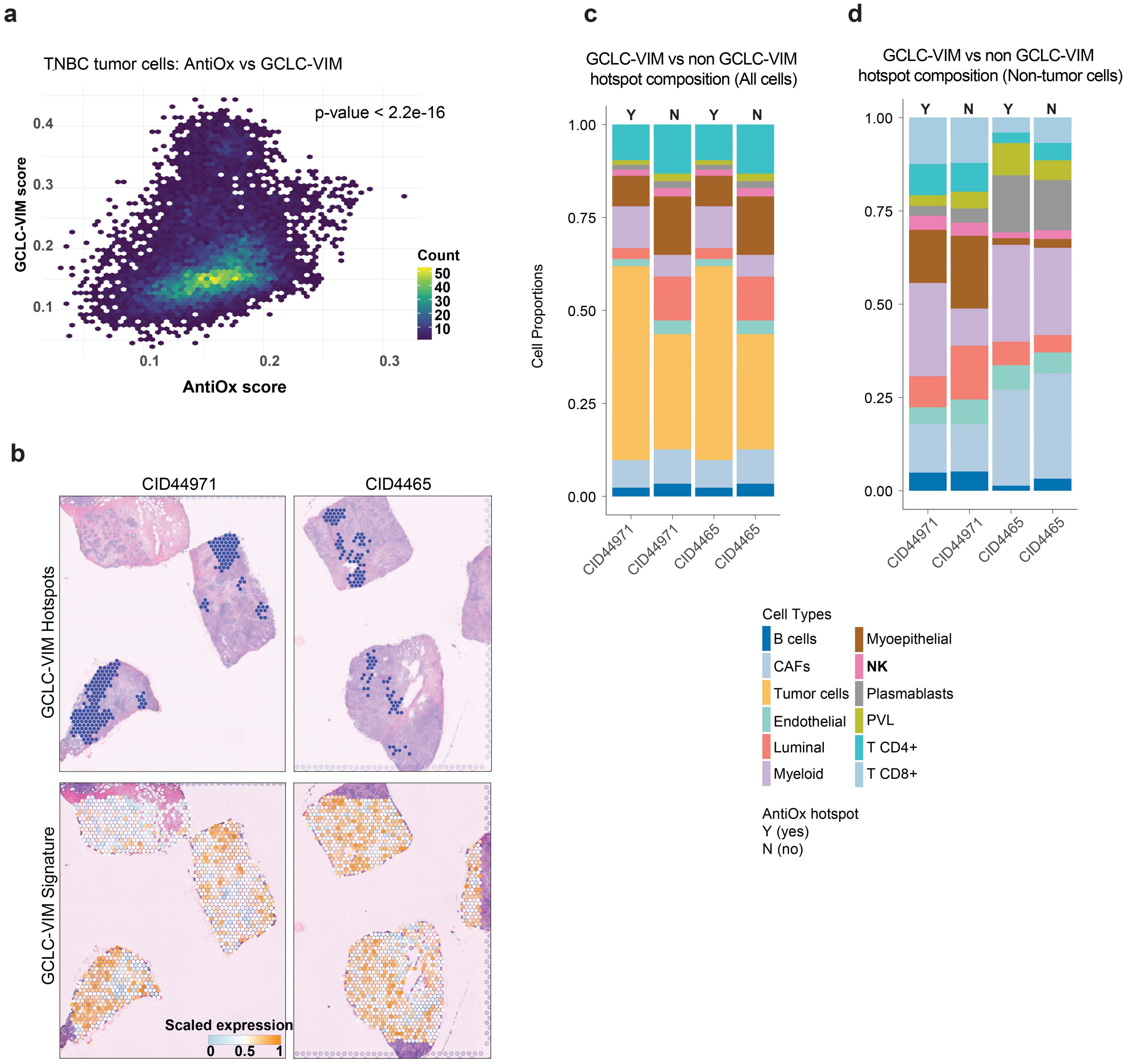

Supplementary Figure 4. **a** A correlation plot between the AntiOx and GCLC-VIM signatures across TNBC tumor cells. These results are from reanalyzed scRNAseq data[19]. **b** H&E Visium data is presented together, colored with the identified GCLC-VIM hot spots (top) and the GCLC-VIM signature expression (bottom) for samples CID44971 (left) and CID4465 (right), respectively. **c-d** Stacked bar plots summarize cell type composition differences in GCLC+VIM+ hot spots and non-hotspots considering all cells (**c**) and rescaled without tumor cells (**d**).
